# Predicting *Alu* exonization in the human genome with a deep learning model

**DOI:** 10.1101/2024.01.03.574099

**Authors:** Zitong He, Ou Chen, Noelani Phillips, Giulia Irene Maria Pasquesi, Sarven Sabunciyan, Liliana Florea

**Author notes:** Corresponding author: Liliana Florea, Ph.D., 733 N Broadway, MRB 453, Department of Genetic Medicine, Johns Hopkins School of Medicine, Baltimore, MD 21205.

## Abstract

*Alu* exonization, or the recruitment of intronic *Alu* elements into gene sequences, has contributed to functional diversification; however, its extent and the ways in which it influences gene regulation are not fully understood. We developed an unbiased approach to predict *Alu* exonization events from genomic sequences implemented in a deep learning model, eXAlu, that overcomes the limitations of tissue or condition specificity and the computational burden of RNA-seq analysis. The model captures previously reported characteristics of exonized *Alu* sequences and can predict sequence elements important for *Alu* exonization. Using eXAlu, we estimate the number of *Alu* elements in the human genome undergoing exonization to be between 55-110K, 11-21 fold more than represented in the GENCODE gene database. Using RT-PCR we were able to validate selected predicted *Alu* exonization events, supporting the accuracy of our method. Lastly, we highlight a potential application of our method to identify polymorphic *Alu* insertion exonizations in individuals and in the population from whole genome sequencing data.

## Introduction

*Alu* sequences represent an order of transposable elements termed Short Interspersed Nuclear Elements (SINEs) that are present exclusively in primates. The three *Alu* families, *Alu*J, *Alu*S and *Alu*Y, have been active and duplicating at different times during evolution, contributing to genomic structural diversity (1–3). *Alu* sequences represent 11% of the human genome, with over one million copies located primarily in introns and intergenic space, and only a small fraction residing within gene exons. *Alu* element insertions into genes may disrupt or create new function, and may affect RNA polyadenylation, editing and splicing (4–7). In particular, *Alu* exonization, in which an intronic *Alu* element is spliced in as a new exon into a transcript, can have drastic consequences for the gene’s function, leading to the production of new protein isoforms or altering the gene’s regulation by introducing a premature termination codon (PTC) that triggers the nonsense mediated decay (NMD) mechanism to degrade aberrant transcripts (8). Through alterations to the gene regulation and function during evolution, *Alu* exonizations have acted to increase functional diversity.

However, they can also have deleterious effects, with cryptic *Alu* exon activation contributing to a number of human genetic diseases (referenced in (6)). While the importance of *Alu* exonization for gene function and regulation is well recognized, its extent and the biological processes in which it is involved are not yet fully understood.

Exonization of *Alu* elements is frequent during primate and human evolution, with thousands of human genes expressing *Alu* exons (8,9). The consensus *Alu* sequence is ∼280 bases in length, formed from two diverged dimers (‘arms’), ancestrally derived from the 7SL RNA gene, separated by a short ‘A’ rich segment. Most exons are generated from the right arm, albeit left arm exonization is also frequent (10), and *Alu* elements are predominantly incorporated in the antisense to the gene (11). The *Alu* sequence contains multiple sites resembling consensus splice signals and other splicing regulatory elements. *Alu* exon activation occurs through mutations, usually at the 5’ end of the *Alu* and in the surrounding introns, that create new and typically weak splice junctions and splicing regulatory elements. In most cases, *Alu* elements are incorporated as alternatively spliced exons, with the new *Alu*-containing transcript becoming a minor isoform (9). In time, *Alu* exons may mutate and acquire new function, with some transcripts eventually becoming the major isoform for the gene (12). A subset of *Alu* exons were shown to have translational potential and produce primate and human specific peptides (13). However, most *Alu* exons surveyed with high throughput sequencing have shown low splicing and expression levels (14). Collectively, these features contribute to the exon recognition by the spliceosome and can also be exploited to design computational prediction models.

*Alu* exonization accounts for an estimated 5% of the alternatively spliced (cassette) exons in the human genome (9). This estimate was based on early analyses of EST and cDNA sequence data and on characteristic features of the exon and surrounding introns incorporated into a machine learning model (15). High throughput sequencing has made it possible to identify *Alu* exons large scale and for a wide variety of conditions, however, significant challenges remain to producing the full human catalog. Processing the massive amount of RNA-seq data would be expensive and would quickly reach diminishing returns.

*Alu*-containing transcripts may further be targeted by NMD, significantly lowering the number of RNA-seq reads and making the events more difficult to detect. Furthermore, some tissue or condition specific events will be missed because source RNA-seq data is not available in sequence databases, while others, such as potentially disease-causing cryptic *Alu* exons, may be unobservable due to constitutive repression by splicing inhibitors such as hnRNPC (16). As a result, *Alu* exons remain under-represented in gene annotation databases.

Lastly, *Alu* retrotransposon insertion into the human genome is ongoing, through germline as well as somatic mutations induced by disease and aging (17). A small fraction of insertions will lead to exonization, contributing to genetic diversity but also to new genetic diseases. Some insertions develop the capacity for exonization, contributing to genetic diversity but also potentially leading to or modulating the severity of disease throughout the lifespan of the individual (18,19). Reported examples of *Alu* exonization events causing disease include a G-to-T transversion activating a cryptic splice site in an antisense *Alu* element in intron V of collagen type IV alpha 3 chain (COL4A3) gene, causing autosomal recessive Alport syndrome (20). Other examples include mutations in the ornithine-delta-aminotransferase (OAT) gene causing gyrate atrophy of the choroid and retina (21); a survivin allele (baculoviral IAP repeat containing 5 (BIRC5)), causing Sly syndrome (22); a fibroblast growth factor receptor 2 (FGFR2) allele causing Apert syndrome (23); and a 6-pyruvoyltetrahydropterin synthase (PTS) allele resulting in tetrahydrobiopterin deficiency (24). In other cases, *Alu* exonization is prone to producing premature termination by frameshifts and may lead to proteins that are deleterious or have modified properties. For example, the interferon alpha and beta receptor subunit 2 gene (IFNAR2) isoform IFNAR2-S, which encodes an *Alu*-derived alternative terminal exon 9b, functions as a decoy receptor that inhibits interferon signaling, including in cells infected with SARS-CoV-2 (25). Nevertheless, the extent of polymorphic *Alu* exonization in the population and its relation to disease is unknown.

To answer these questions, we propose an unbiased approach to detecting *Alu* exonization in the human genome using a deep learning convolutional neural network (CNN) model. To our knowledge, this is *the first* machine learning approach to comprehensively and accurately model *Alu* exonization. The model, implemented in the software eXAlu, learns *Alu* exon information from the genomic sequence alone, without the need for RNA-seq data. eXAlu operates in multiple stages, combining six convolutional and two fully connected layers. Unlike RNA sequencing-based approaches, it analyzes each input *Alu* sequence only once, and therefore is highly efficient. It also eliminates the need for extensive and condition specific RNA sequencing, therefore offering an unbiased approach to discovery. The model’s salient features and implications for science include:

- High accuracy (Sn=0.88, Pr=0.78, AUC=0.89 on the *test* data; AUC=0.91 on the *training* data), making it feasible to apply at genome-scale;
- A training collection of 11,930 distinct exonized *Alu* sequences, extracted from RNA-seq data in 28 human tissues from the GTEx repository (26), twice as large as the number of events (5,076) currently represented in the GENCODE v.36 database;
- Recognition and prediction of sequence elements important for the recognition of exonized *Alu* sequences, likely regulatory elements, as illustrated in an analysis of 870 *Alu* exonization events in human frontal cortex (14);
- A significantly higher estimate of the number of exonized *Alu* elements in the human genome, 55-110K, 11-21 times larger than earlier estimates and their representation in the GENCODE reference gene annotations;
- A potential practical approach to predicting polymorphic *Alu* exonizations in population genotype data, as evidenced by an analysis of 28,319 polymorphic *Alu* insertions previously reported by the 1000 Genomes Project that produced 97 candidate exonization events, of which 3 could be validated in donor-matched LCL RNA-seq data (27).

The eXAlu program was written in Python using the package PyTorch. Software is available free of charge under a GPL license from https://github.com/splicebox/eXAlu.

## Results

We describe eXAlu, a Convolutional Neural Network (CNN) model that estimates the likelihood for an *Alu* element to undergo exonization based on the *Alu* and surrounding genomic (pre-mRNA) sequence. We show that eXAlu achieves high accuracy on data from multiple human tissues and therefore can be effectively applied genome-scale. We then apply eXAlu to a set of previously reported *Alu* exons from human frontal cortex, and demonstrate that it can identify regulatory features important for exonization. Further, by scanning the *Alu* repeat annotations in the entire human genome, we identify ∼63K high confidence *Alu* candidates for exonization, considerably exceeding the current estimate (9) and the current representation in gene annotation databases. We validate several randomly selected predictions with RT-PCR in liver and cerebellum tissue. Lastly, we illustrate a potential application of eXAlu in determining putative exonization events at sites of polymorphic *Alu* insertions directly from whole genome sequencing data, without the need for RNA-seq data, as a first step in creating a catalog of variation in the human population.

### Overview and evaluation of the algorithm

eXAlu predicts the potential for an *Alu* element to be spliced in as an exon based on a deep learning model of the *Alu* and surrounding (350 bp) flanking sequences. The model consists of six convolutional layers, to extract features from the sequences, followed by two fully-connected layers, to calculate the likelihood score. The output is a label indicating whether the *Alu* can be exonized (**Figure 1A** and **Methods**).

**Figure 1.**
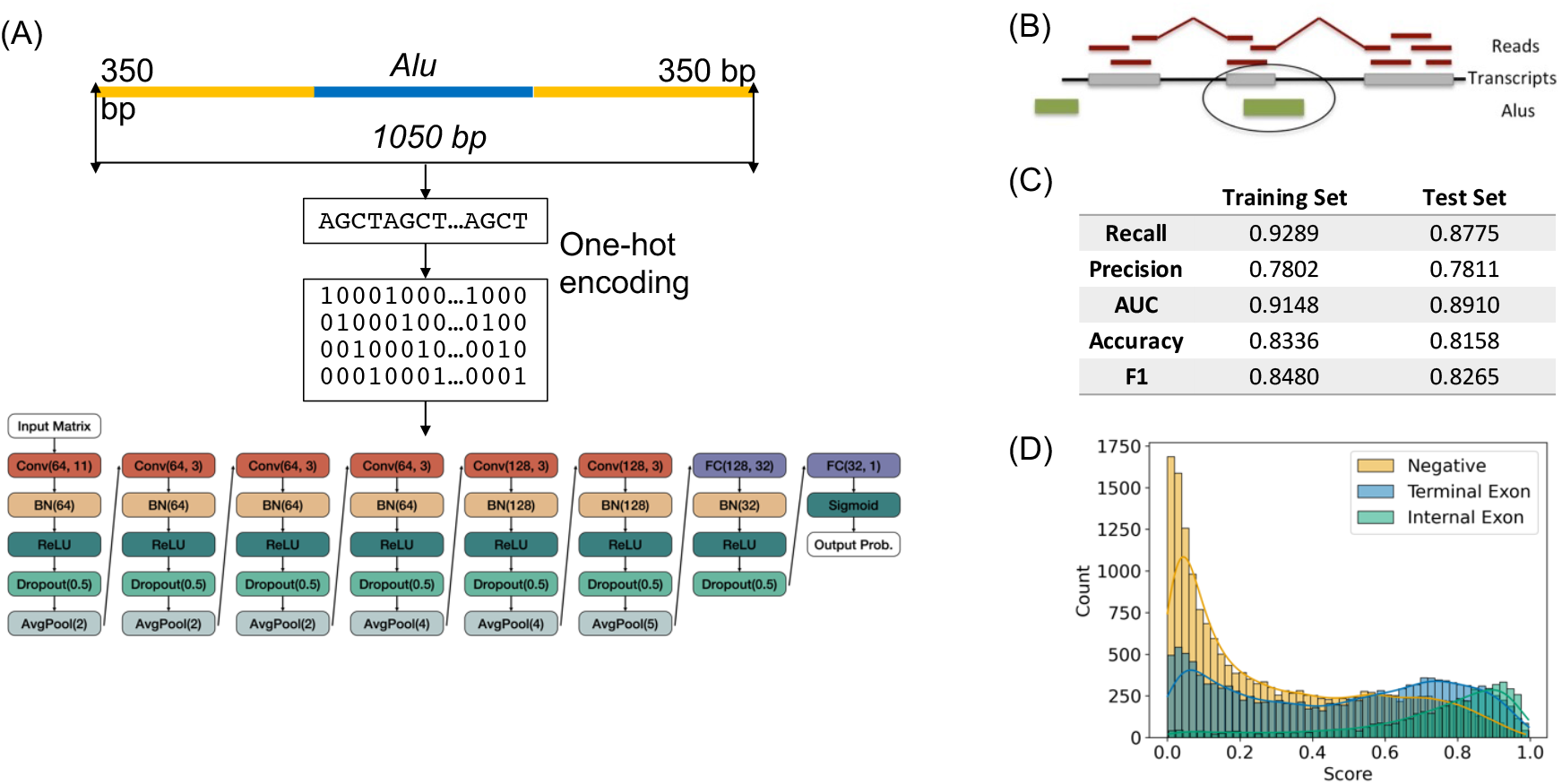
The Deep Learning model of *Alu* exonization. (A) The input consists of *Alu* and 350 bp context sequences, encoded using one-hot encoding. The model consists of a Convolutional Neural Network (CNN) with 6 convolutional layers, to extract features from the sequences, and two fully connected layers, to calculate the likelihood score. The output is the probability score that the input *Alu* is exonized, labeled as follows: the *Alu* is deemed ‘exonized’ iff (score>=0.5). (B) Collection of training data. Aligned RNA-reads are assembled into transcripts, and internal exons 40-400 bp long overlapping antisense *Alu* annotations are selected as *Alu* exons. (C) Performance of the CNN model. (D) Histogram of scores for GENCODE *Alu*-overlapping exons, terminal exons versus internal exons, and comparison to negative (non-GENCODE) examples.

To develop and evaluate the models, we collected 12,206 exonization events at 11,930 *Alu* elements, extracted from RNA-seq data in 276 samples from 28 human tissues obtained from the GTEx project (**Figure 1B**, **Supplementary Table S1**, and **Methods**). Briefly, following read mapping and transcript assembly, 40--400 bp internal exons overlapping an annotated *Alu* element located on the opposite strand were classified and retained as *Alu* exons. (While exonization can occur in either direction, it is reported to strongly favor the reverse orientation (6,11)). We split the data into non-overlapping training and testing sets, with a 8:1 ratio.

Following calibration, the model achieved high performance (AUC=0.89-0.91, Acc=0.82-0.83 F1=0.83-0.85) on the training and the test sets (**Figure 1C**). Sensitivity was Sn=0.88 and 0.93, respectively, for the two sets, and precision Pr=0.78 for both, indicating that eXAlu is suitable to apply to genome-wide data. Further, program sensitivity on two data sets generated from two previously unused tissues, salivary gland and uterus, and on *Alu* elements not included in the training set, representing 676 and 764 *Alu* sequences, was 0.89 and 0.90 respectively, indicating that the program learned sequence features predictive of the exonization potential and can be used for general data sets.

We also tested the ability of the model to distinguish between *Alu* elements embedded in the gene 5’ and 3’ UTR sequences versus those recruited as internal exons (‘exonized’). The two classes showed significantly different score distributions, pointing to different adaptations to selective pressures, which were also distinct from those of *Alu* elements not included in gene bodies (**Figure 1D**).

### Features of *Alu* exons in human frontal cortex data

To understand the power and any limitations of the model, and to identify characteristic features of the *Alu* and context sequences, we applied eXAlu to a well characterized data set. We previously reported 1,019 exonization events at 870 *Alu* loci in human frontal cortex extracted from RNA-seq data in 117 GTEx samples, of which 832 *Alu* elements were between 100 and 350 bp and could be used for our study. The sequences contained elements in all three families (*Alu*J, *Alu*S and *Alu*Y) and therefore likely underwent exonization at different timepoints, were predominantly alternatively spliced, and were generally included at low splicing and expression levels (14). eXAlu correctly predicted 779 sequences and missed 53, for a sensitivity of 0.94.

#### Reported features of exonized Alu sequences

While deep learning methods do not rely on an *a priori* identified set of features, earlier studies have assayed features associated with exonized *Alu* sequences, including several among: *Alu* arm, *Alu* length, length of *Alu* exon, lengths of surrounding introns, *Alu* positioning relative to the splice sites, splice signal strength and stability of the *Alu* secondary structure (15). We *first* assess the features comparatively between the true positives (TP) and false negatives (FN) in the frontal cortex data, to assess biases in the model and to identify potential areas for future improvement. We observed *no* statistically significant correlation between true positive and false negative predictions and originating *Alu* arm (Х^2^-square test, p-val=0.056), *Alu* length (Kolmogorov-Smirnov test, p-val=0.150), exon length (Kolmogorov-Smirnov test, p-val=0.139) or the lengths of flanking introns (Kolmogorov-Smirnov test, p-val=0.692), or distance between the *Alu* exon splice sites (donor, acceptor) and *Alu* endpoints (Kolmogorov-Smirnov tests, p-val=0.109 and 0.852). The only feature that showed a significant positive association was the donor splice signal strength (Х^2^-square test; p-val=5.3e-5), whereas the acceptor splice site strength was uncorrelated (Х^2^-square test, p-val=0.146), indicating that a strong 5’ but not 3’ intron end is important for recognition and potentially also for exonization (**Figure 2A**). Similarly, there is no statistically significant difference between the stability of the *Alu* RNA secondary structure between the two classes (Kolmogorov-Smirnov, p-val=0.345), albeit the per base minimum free energy (MFE) of false negative examples appears generally higher (lower stability) compared to the predicted examples. Notably, previous studies have focused exclusively on shorter exons that involve a single *Alu* arm (10,15), primarily the right arm. Therefore our method, which achieves 0.91 sensitivity (222 out of 250) in recognizing exonization events spanning both *Alu* arms, brings unique capabilities as well as insights into this new class and potential new mechanism of exonization.

**Figure 2.**
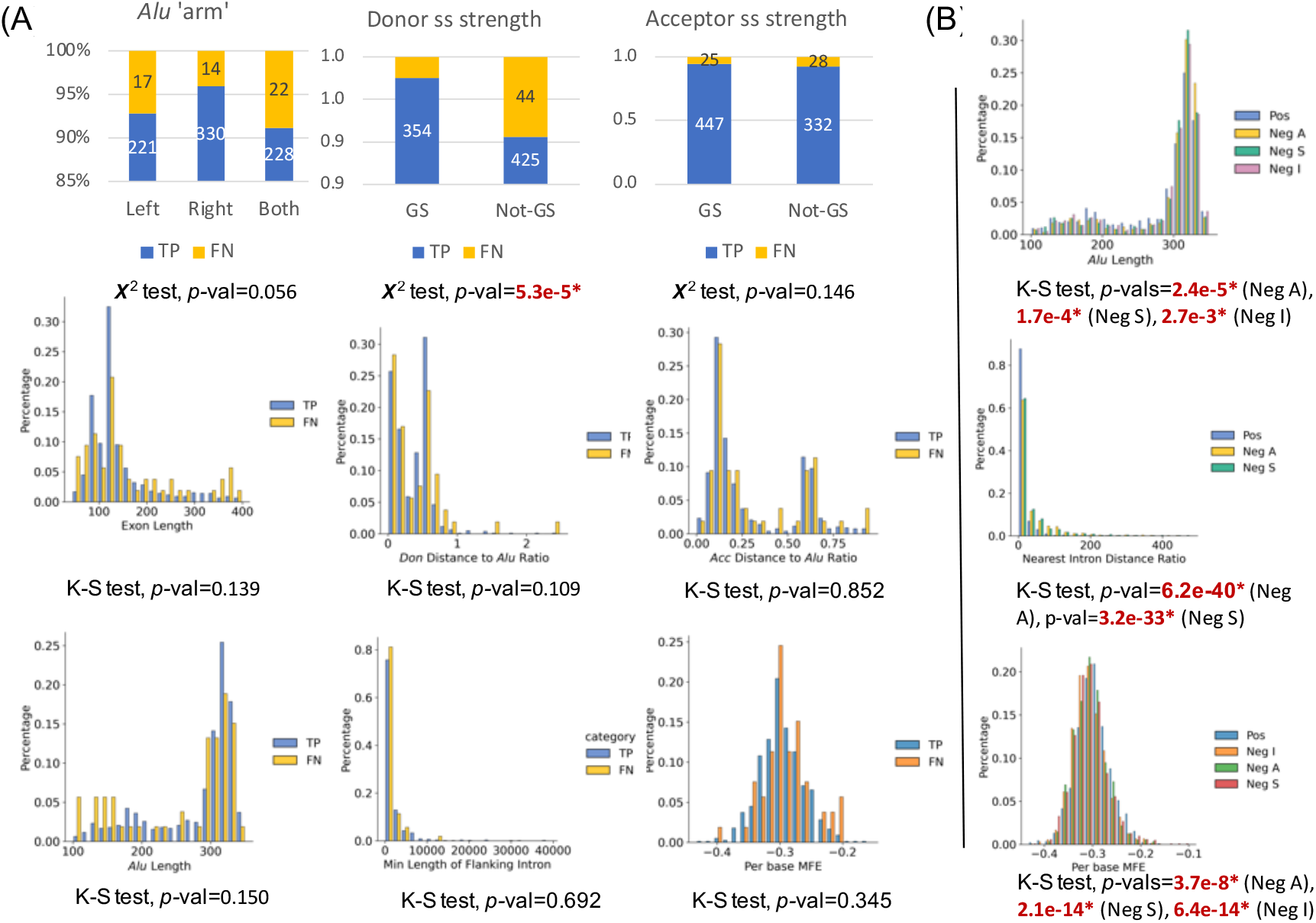
Assessment of previously reported features predictive of *Alu* exonization. (A) Correlations of *Alu* arm, donor and acceptor site strength, exon length, splice site to *Alu* endpoint distance, *Alu* length, length of flanking introns, and stability of the *Alu* RNA secondary structure, between true positive (predicted) and false negative (missed) examples. (B) Correlations of *Alu* length, flanking intron length and stability of the *Alu* RNA secondary structure, with the ‘positive’ (known exonized *Alu* elements) and ‘negative’ (*Alu* elements not known to be exonized) categories. ‘Neg S’ - intronic and on the same strand as the gene (‘sense’); ‘Neg S’ - intronic and on the opposite strand to the gene (‘antisense’); and ‘Neg I’ - intergenic.

*Next*, to identify characteristics that distinguish exonized from non-exonized *Alu* sequences, we compare features between all frontal cortex (‘positive’) and matched ‘negative’ (control) examples, to revisit previous findings. We consider three control classes: intronic *Alu* on the opposite strand to the gene that were *not* included in our training set (‘Neg A’), intronic *Alu* on the same strand as the gene (‘Neg S’), and intergenic *Alu* elements (‘Neg I’). (As a caveat, we note that any of these datasets may contain examples that are in fact exonized but whose status at the time of this selection is unknown.) There was a statistically significant association with the length of the *Alu* sequence (Kolmogorov-Smirnov test, p-values 2.4e-5, 1.7e-4 and 2.7e-3 for the three categories), exonized *Alu* repeats being statistically shorter; with the distance to the nearest exon (Kolmogorov-Smirnov test, *p-*values 6.2e-40 and 3.2e-33, for the ‘Neg A’ and ‘Neg S’ categories), exonized *Alu* elements being more likely to be located in the vicinity of an existing exon; and, lastly, with the stability of the secondary structure, exonized *Alu* elements having a less stable RNA secondary structure compared to their non-exonized counterparts (Kolmogorov-Smirnov test, *p-*values 3.7e-8, 2.1e-14 and 6.4e-14 for the ‘Neg S’, ‘Neg A’ and ‘Neg I’ categories, respectively) (**Figure 2B**).

#### Inferred sequence features important for Alu exonization

To determine features important for model recognition, and presumably for the exonization process, we performed an *in silico* mutagenesis experiment. For each of the 832 exonized *Alu* elements, we determined the effect of all possible single nucleotide sequence mutations (SNPs) in the *Alu* sequence and its surrounding regions (350 bp) and plotted it along the genomic interval. Negative peaks (see **Methods**) then point to changes in the local sequence context that reduce the probability of exonization, and positive peaks mark changes leading to an increased probability. Negative peaks were predominantly present around splice sites, but also within 20-50 bp proximity of splice sites within the exon or in the intron, reflecting the model’s learning of splice site motifs and other sequence features, potentially binding sites of splicing-related RNA binding proteins. (In contrast, there were few positive peaks.) Three examples of regions with peak and splice site annotations, at the BCL7B, CCDC171 and the RPP30 genes, are shown in **Figure 3A**. We searched two of the peak sequences, at the CCDC171 and the RPP30 genes, against the AtTRACT database (28), which contains experimentally validated profiles of known RNA binding proteins (**Figure 3B**). The search for the first motif, CAGCCUCC, located 17 bp downstream of the acceptor splice site within the *Alu* exon (chr9:15888983-15889101), identified putative binding sites for the KH-Type Splicing Regulatory Protein (KHSRP) factor, a multifunctional RNA-binding protein implicated in a variety of cellular processes, including transcription, alternative pre-mRNA splicing, and mRNA localization (29,30). Similarly, searching for the AUUUUUG motif, located 34 bp upstream of the acceptor site in the intronic region, retrieved a potential binding site for the heterogeneous nuclear ribonucleoprotein C (hnRNPC), indicating possible repression of the exon (16). Therefore, our model learns and can retrieve splicing regulatory features important for *Alu* exonization.

**Figure 3.**
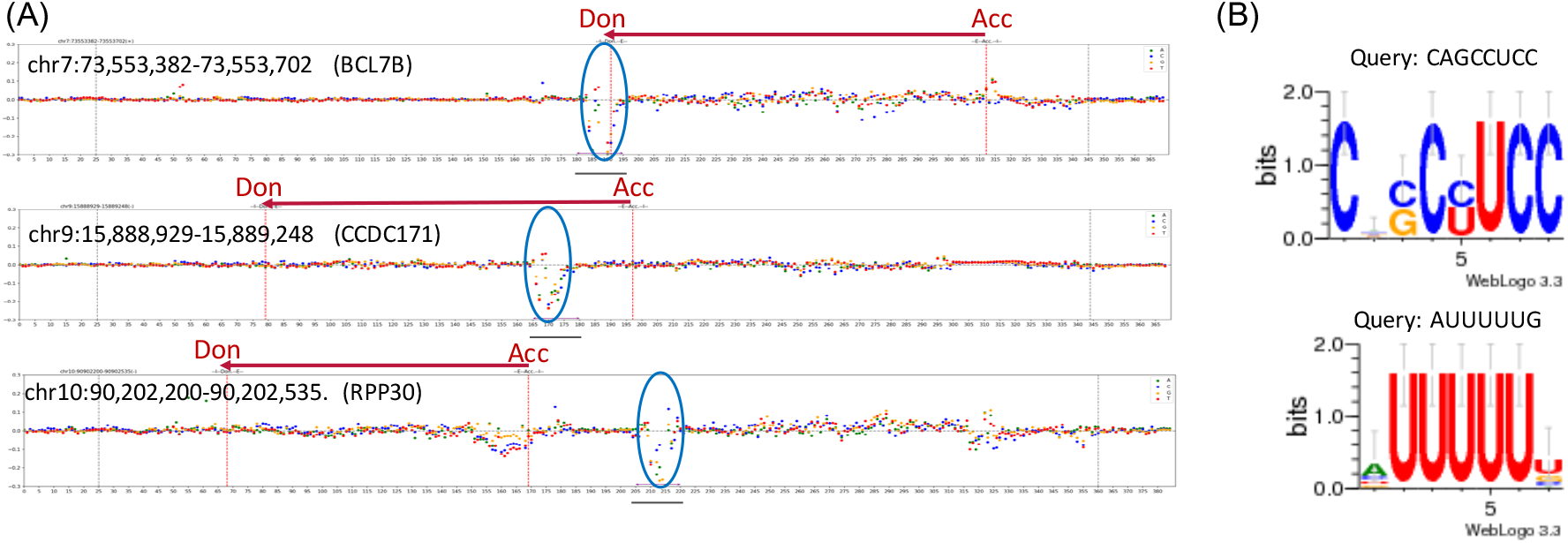
Inferred sequence features important for exonization. (A) Mutagenesis plots at the BCL7B, CCDC171 and RPP30 gene loci exhibiting *Alu* exonization. Plots show the effects on the *Alu* score when each base is individually mutated to each of the three alternate nucleotides (A – green, C – blue, G – orange, T – red) along the genomic interval containing the *Alu* sequence and the 350 bp flanking regions. Only values between –0.3 and 0.3 (Y-axis) are shown. *Alu* endpoints are shown with black vertical bars, exons boundaries with red bars, and peaks are marked with a horizontal bar at the bottom of the plots and highlighted with blue oval marks. (B) Examples of splicing regulatory motifs inferred from the differential score peaks.

### Estimating the extent of *Alu* exonization in the reference human genome

We applied the eXAlu model to the 1,091,952 *Alu* sequences between 100-350 bp annotated in the reference human genome GRCh38. Of these, 140,149 elements were located within gene regions, predicted to be exonized, intronic, and in antisense to the gene. When searching for evidence of exonization in the input data sets, 62,781 (44.8%) elements had evidence in at least one of the categories: GENCODE *Alu* exons (4,318), *Alu* exons represented in the RefSeq or UCSC reference gene annotations (1,653), *Alu* repeats in the training set (8,427), and splice junction support from spliced read alignments in the collected tissue RNA-seq data sets.

Because the GRCh38 assembly still has hundreds of gaps, including at repetitive regions such as the centromere and the short arms of acrocentric chromosomes, to further assess the potential for *Alu* repeat exonization in a complete genome we applied our model to the 1,204,971 qualified *Alu* annotations in the CHM13 v2.0 assembly (31). CHM13 represents the first truly complete genome of a single individual, and in which all gaps have been filled. We then projected and compared the sets of exonized *Alu* elements as predicted by eXAlu and located within the introns and in antisense to annotated genes, herein called exonized ‘core’, between the two genomes. While the size of the ‘core’ is comparable between the two assemblies, with more than 80% in common, more exonized *Alu* elements in the GRCh38 genome are located outside of a gene’s context in CHM13, likely because the gene annotations of the latter are less complete (**Table 1**). Intriguingly, more GRCh38 ‘core’ elements have counterparts that are not predicted to be exonized in the CHM13 genome, potentially due to their sequence context. Lastly, there are 883 CHM13 ‘core’ elements that could not be projected onto the hg38 genome, representing either polymorphisms or repeats in gapped areas of the GRCh38 genome. Overall, the exonized *Alu* content in the two genomes appears to be largely concordant, with more investigation needed to reveal the nature of their differences.

**Table 1.** Comparison of exonized ‘core’ and *Alu* repeat universe between the GRCh38 and the CHM13 human genome assemblies.

|  | **GRCh38** | **CHM13** |
| --- | --- | --- |
| Exonized ‘core’ size | 140,155 | 136,477 |
| **Status in the other genome** |  |  |
| Exonized ‘core’ | 113,763 | 113,603 |
| Exonized, not in gene context | 16,655 | 9,231 |
| Not exonized | 8,800 | 4,337 |
| Unmatched | 849 | 8,393 |
| Not transferred | 88 | 883 |

Lastly, to more comprehensively and reliably assess the confidence of our predictions, we randomly selected ten novel candidate events (nine antisense intronic and one intergenic) out of the 16,380 predictions on chromosome 12 from among those with tissue splice evidence, for experimental validation with RT-PCR (**Figure 4, Supplementary Table S2** and **Supplementary Figure S1**). At pre-design time, two examples (at the genes AC068987.2 and TTC41P) had incomplete splicing patterns and were excluded. Of the remaining eight cases, RT-PCR testing in cerebellum and liver tissue validated both junctions for four examples (at the genes RESF1, DIP2B, AACS and KDM2B), one junction for two examples (chr12:57,259,972-57,260,295, at gene R3HDM2, and chr12:121,782,877-121,782,881, at gene RHOF – data not shown), and primer design failed for two examples, at the PITPNM2 gene and at the intergenic locus (**Figure 4**). Given the technical challenges associated with specifically amplifying repetitive regions, our ability to validate six out of the eight events that could be tested by PCR, and in effect six out of the seven that fulfill our criteria for eligibility (intronic and in antisense to the gene), supports the validity of our *Alu* exonization prediction tool.

**Figure 4.**
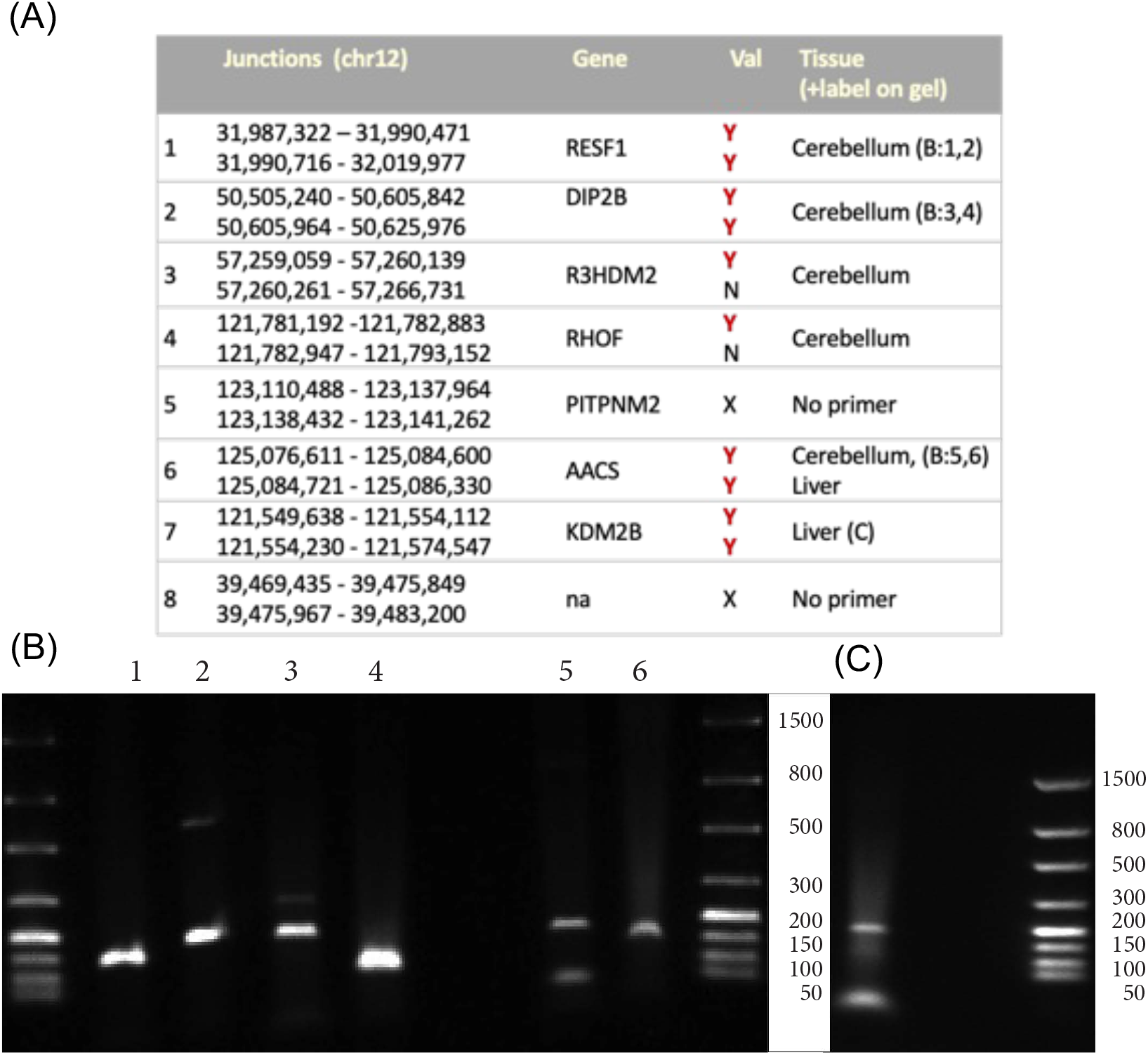
PCR validation of selected *Alu* exons. (A) Summary of PCR validation experiments. (B) For three *Alu* elements, both the 5’ and the 3’ splice junctions between the *Alu* and the adjacent (flanking) exons were verified by PCR. The column numbers (**1,2**) correspond to chr12:31990430-31990748 (RESF1), (**3,4**) to chr12:50,605,807-50,606,132 (DIP2B), (**5,6**) correspond to chr12:125,084,561-125,084,891 (AACS). (C) PCR amplification of the adjoining exons that include the spliced *Alu* element for chr12:121,554,076-121,554,260 (KDM2B).

### Prediction of *Alu* insertion polymorphisms undergoing exonization in individual genomes

*Alu* retrotransposons continue to insert in the genomes of individuals, with some instances further undergoing polymorphic exonization. To explore this hypothesis and the potential of our method to identify such instances, we applied eXAlu to a set of 28,319 polymorphic *Alu*Y insertions previously discovered from whole genome sequencing data in 2,504 participants in the 1000 Genomes Project (27). We retained 8,909 variants that occurred within introns and in antisense to the annotated genes, which were analyzed with eXAlu. eXAlu predicted 213 *Alu* sequences (score>=0.5) potentially exonizable. We next searched for evidence of exonization of each of these insertions in the genome of origin, using matched RNA-seq data from Lymphoblastoid Cell Lines (LCLs) from the same individual, where available, sequenced as part of the GEUVADIS project. While RNA-seq data were only available for a fraction of participants (465 out of 2,504), 97 of the *Alu* polymorphic insertions predicted to be exonized had matched LCL RNA-seq data, in a total of 320 RNA-seq samples. Of those, we manually curated and reconstructed the exon-intron structure of 3 examples (**Figure 5** and **Supplementary Figure S2**), at the Transcriptional Coactivator and Phosphatase 3 (EYA3), 3-Hydroxy-3-Methylglutaryl-CoA Reductase (HMGCR) and Dehydrogenase/Reductase X-Linked (DHRSX) genes, using RNA-seq evidence from spliced alignments flanking the *Alu* insertion. While the validated number of predicted events is small, being severely limited by the fact that RNA-seq data is only available for a subset of individuals and from a single tissue, these examples highlight the potential of our model to identify instances of polymorphic *Alu* exonization in population genomic data.

**Figure 5.**
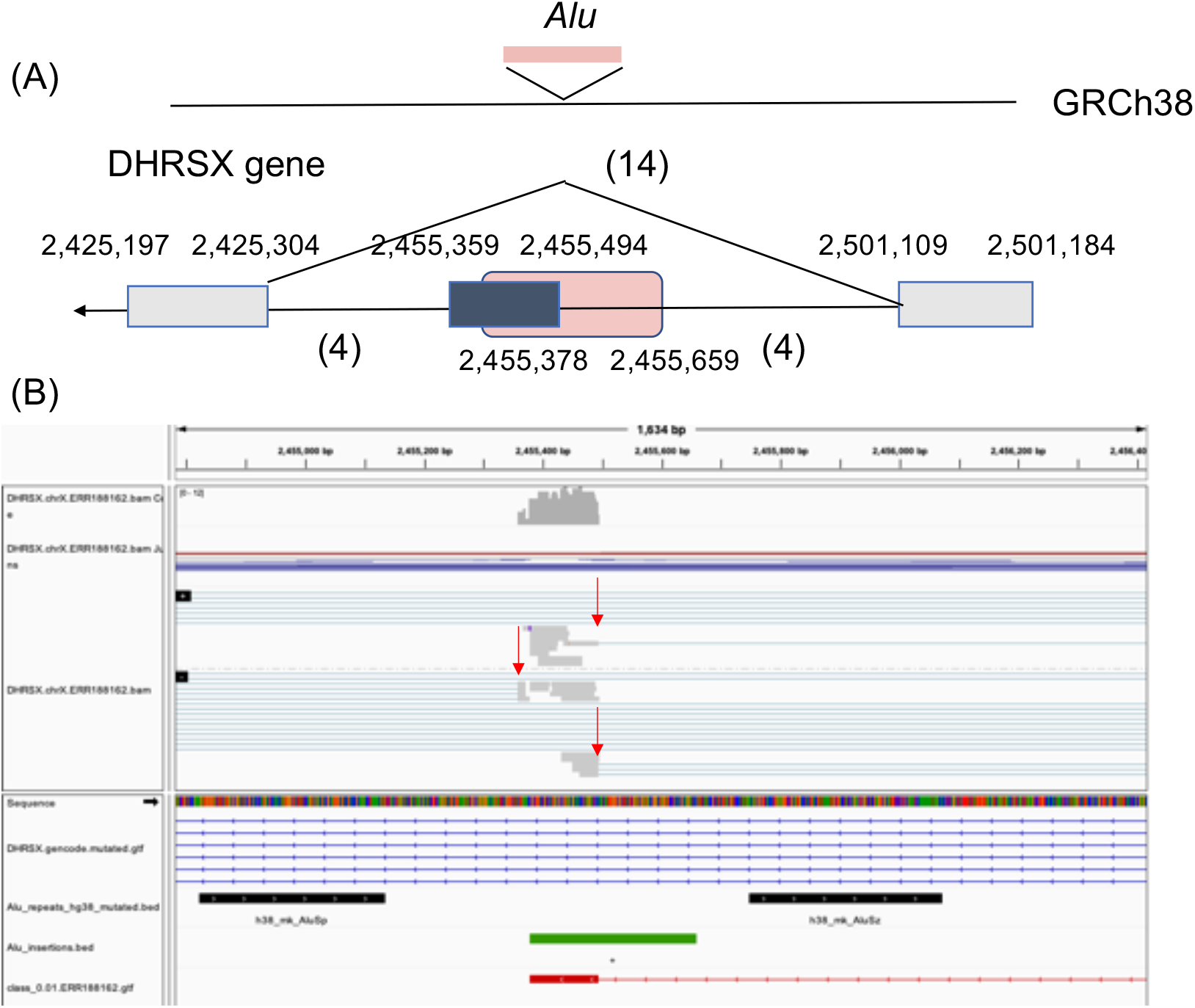
Example of polymorphic *Alu* insertion at the DHRSX gene locus predicted to be exonized, and GEUVADIS RNA-seq evidence. (A) Gene structure diagram showing exonization of the inserted *Alu.* The *Alu* element is shown in pink, and exons in grey boxes. Numbers in parenthesis represent the number of RNA-seq spliced alignments in support of the splice junction. (B) IGV representation of spliced alignments demarcating the *Alu* exon (grey, top panel), fixed (in the hg38 genome) *Alu r*epeats (black horizontal bars), polymorphic inserted *Alu* repeat (green), and transcript assembled from the RNA-seq reads showing the Alu exon boundary (red), all shown along the mutated genome.

### Discussion and Conclusions

The extent and role of *Alu* exonization in primate evolution and in human disease are not yet fully understood. While *Alu* exons were initially estimated at 5% of the alternatively spliced (cassette) human exons, recent studies using deep RNA sequencing technologies have hinted at a potentially more widespread presentation (32). While some *Alu* exonization events affect gene function directly, through changes to the protein (13), it is anticipated that many *Alu* containing transcripts contribute to regulating gene expression by modulating the relative expression of isoforms through degradation by nonsense mediated decay (NMD). A number of events are conserved across primates, raising the question of an evolutionary gene regulatory mechanism (25). Therefore, characterizing *Alu* exonization in a comprehensive way would allow not only for in depth functional studies at the gene level but also for mechanistic studies, as the events can be observed in other species and in model organisms.

Cataloging *Alu* exonization events from RNA sequencing data has been fruitful but is computationally expensive and suffers from diminishing returns, with some condition-specific events potentially never to be identified. We developed a machine learning model to determine high likelihood *Alu* exonization events without the need for RNA sequencing data, efficiently and accurately. Deep learning models, in particular CNNs, have been successfully employed in bioinformatics applications in recent years, and have the advantage of circumventing the need for *a priori* determining an informative set of features. Traditionally, deep neural networks require large amounts of training data; however, eXAlu takes advantage of the high similarity between the sets of positive and negative sequences, such that less data is needed to identify only distinguishing features. As training set, we collected 11,930 *Alu* exonization events from multiple human tissues, which represents the largest such collection to date, twice as large as the number of events represented in the GENCODE annotations (5,076). The data set is diverse across *Alu* families, and includes *Alu* elements beyond those originally reported, including both-arms exonizations. An early machine learning (SVM) method (15) based on a set of distinguishing features was reported to achieve high accuracy, however, it was based on a limited set of 390 elements, exclusively originating from the left or the right (but not both) arm of the *Alu*. eXAlu is able to predict cross-arm exonization instances, and therefore can shed light on this type of exonization.

Our method achieves up to 0.93 sensitivity and 0.78 precision. One limitation to performance may be the presence of exonizable elements among those used as our set of negative examples, affecting both the training and the performance measurements, and which can be improved with better and more complete gene annotations. Our method could also be improved by incorporating additional distinguishing features. In particular, the stability of the local RNA secondary structure was identified as a discriminating feature of exonized *Alu* elements, potentially due to increased accessibility to splicing and other regulatory factors (15,33,34). It was also a distinguishing factor in our analyses comparing positive and negative examples, and predicted versus missed events in the frontal cortex data. While our model may reflect characteristics of RNA secondary structure, the linearity of the CNNs does not naturally render them as optimal models. Explicitly incorporating features of secondary structure could therefore prove productive.

Our program predicts 140,149 *Alu* elements in the human genome located in introns and in antisense to the gene that have the potential to undergo exonization, of which ∼63,000 have evidence in the form of GENCODE annotations or GTEx RNA-seq data. We additionally obtained a high validation rate with RT-PCR for a limited number of randomly selected candidates. Therefore, given our program’s estimated performance metrics, we extrapolate that between 55-110K *Alu* elements in the human genome with these characteristics (*i.e.*, intronic and in antisense) may be undergoing exonization in the human genome, or 11--21 times more than the 5,076 events currently annotated in GENCODE v.36. This widespread presentation of *Alu* exonization events in the human genome renews questions about their role in gene regulation and function.

Lastly, we reviewed a potential application of our tool in detecting human genetic variation, to predict likely exonization events at loci of polymorphic *Alu* insertion. Our experiment based on the GEUVADIS data was severely limited by the number of samples, the availability of genotype and RNA-seq data (for validation) from the same individual, and the fact that RNA data from only one tissue was available, whereas *Alu* exons may be selectively expressed in tissues and cell types. Nevertheless, the identification of three such events supported by RNA-seq evidence is promising, indicating that the current tool can be further optimized for use in identifying potentially disease-causing rare and polymorphic *Alu* exonization events from population genotype data.

## Methods

### The *Alu* exonization CNN model

Our Deep Learning model predicts the likelihood that an *Alu* element will become exonized. Implicitly, the model tests the hypothesis that the genomic context consisting of the *Alu* and short flanking sequences is sufficient for exonization. The input for the model are 400 bp sequences centered on the *Alu* sequence and 25 bp surrounding regions, ‘N’-padded if necessary. Only *Alu* sequences between 100-350 bp are used. The output label is a boolean, namely 1 (positive) if the *Alu* is predicted to cause exonization (score>0.5), and 0 (negative) otherwise (score<=0.5). We use one-hot encoding to map a DNA or RNA sequence (a 1 × N vector) to a 4 × N matrix (**Figure 1A**).

The deep neural network architecture (**Figure 1A**) contains six convolutional layers, Conv(N, S), where N denotes the number of convolutional kernels and S denotes the size of each kernel, to extract features of the input sequence. The Conv modules are followed by batch normalization BatchNorm(N) modules and a average-pooling layer AvgPool(P), where P is the kernel size. The last component of the model is a group of fully connected layers FC(I, O), where I and O are the input and output sizes. We used the Rectified Linear Unit (ReLU) as activation function for the convolutional layers and the first fully connected layer. The second FC uses Sigmoid S(x) to map the network’s output logits from [−1, 1] to [0, 1] classification scores. Additionally, to reduce overfitting, we use Dropout(D) (35) with a rate D of 0.5 for convolutional layers and the first fully connected layer.

To train the model, we use binary cross-entropy LBCE as the loss function,

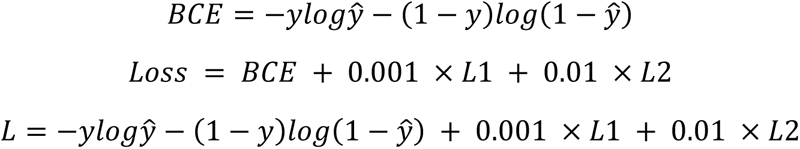

where L1 and L2 denote the L1- and L2-normalizations, and y is a binary label: y = 1 (positive) or y = 0 (negative). Based on the properties of the sigmoid function and binary cross-entropy, we select 0.5 as threshold for predicting a positive (>= 0.5) or negative (< 0.5) value.

### Model training and evaluation

#### Training data

To train and evaluate the model, we extracted *Alu* exonization events from RNA-seq data from 28 human tissues obtained from the Genotype Tissue Expression (GTEx) project. Briefly, reads were mapped to the human genome GRCh38 with the software STAR v2.4.2a (36) and assembled into transcripts with the tool CLASS2 v.2.1.7 (37). Internal exons 40-400 bp long overlapping an *Alu* element located on the strand opposite to the gene were selected as candidates for *Alu* exonization events.

#### Model training and calibration

We randomly divided the data into disjoint testing and training sets, in a 1:8 ratio. We employed the conventional Accuracy = (TP+TN)/(TP+TN+FP+FN), F1 = 2 *Sn*Pr/(Sn+Pr), where Sn = TP/(TP+FN) and Pr=TP/(TP+FP), and the area under the curve (AUC), as performance measures. The AUC is equal to the probability that a classifier will rank a randomly chosen positive instance higher than a randomly chosen negative one. We trained the model for 140 epochs with a learning rate of 0.0005, using Adam (38) optimization. The batch size was set to 2048. In the end, we chose the combination of parameters that maximized the F1-value.

### Analysis of frontal cortex data

#### In silico saturation mutagenesis

For each input *Alu* and surrounding context sequence, we mutated each base (independently) into each of the three other bases, plotting the difference in the model’s score: *ΔS = S(mut)-S(orig)*. To identify areas of consistent change in score, which could mark sites important for *Alu* exonization, we use a sliding window protocol to scan the plot and identify ‘peaks’, either positive or negative. The program uses two sliding windows, namely a ‘core’ window (size = 10, step = 5) to detect candidate peaks, and a ‘vicinity’ window (size = 50, step = 5), to filter false local peaks in an area of high variability. Within each window, the variance (Var *= σ^2^/μ*) of the effect score *ΔS* is calculated over all positions within the window, for all base substitutions and the original sequence. First, a candidate peak is detected when the variance in the ‘core’ window exceeds 2.2 fold the global variance: *Var_c_(i) > 2.2 Var_g_.* Then, the variance in the ‘vicinity’ window, Var_n_, is used to filter ‘local’ likely artifactual peaks in areas of high variability. A ‘core’ peak passes the filter if *Var_c_(i) > 2 Var_n_(i)*. Consecutive peaks are merged for reporting. Parameters including window size, step and filter cutoffs were determined by empirical testing.

#### Prediction of regulatory elements

To determine potential regulatory factors, we searched the sequences of two positive peaks, located within exonic and intronic sequence respectively, against the AtTRACT database, containing Position Weight Matrix representations of motifs of RNA binding proteins, using the ‘Scan sequence’ tool with the default parameters to determine potential binding motifs and selecting the top candidates based on the Exon250 score.

#### Analyses of reported features of Alu exonization

We assessed reported features of *Alu* exonizations to characterize the set of predictions for the 832 qualified (between 100-350 bp long) exonized *Alu* sequence in the frontal cortex data set. First, we tested for features differentially represented among the TP and FN subsets, which could underline biases and limitations in the current model. The following features were considered: i) *Alu* length was determined from RepeatMasker (39) annotations obtained from the UCSC Genome Browser; ii) exon length was obtained from the assembled transcripts, as described above; iii) *Alu* arm was determined based on overlap (>=15 bp) with the 5’ and 3’ halves of the *Alu* sequence; iv) ‘strong’ splice site - splice signal strength was calculated with GeneSplicer (40), separately for the acceptor and donor sites, and sites reported by the software as ‘Medium’ or ‘High’ were counted as ‘strong’; v) distance from splice site to *Alu* endpoint (acceptor site to *Alu* 5’ end, and *Alu* 3’ end to donor site); and vi) the stability of the secondary structure assessed via the Minimum Free Energy (MFE), determined with the RNAfold program from the Vienna v2.6.3 package (41) (the lower the MFE, *i.e.* the more negative, then the more stable the RNA secondary structure). Second, we assessed features including the presence of predicted ‘strong’ splice sites, as reported by GeneSplicer, distance of *Alu* sequence to the nearest exon, and stabiity of the *Alu* RNA secondary structure, for their ability to distinguish between exonized *Alu* sequences (‘positive’) and *Alu* sequences not known to be exonized (‘negative’). Three ‘negative’ data sets were considered: a) intronic *Alu* sequences on antisense strand and not in the input data set (‘Neg A’); b) intronic *Alu* sequences on the gene strand (‘Neg S’); and iii) intergenic *Alu* repeats (‘Neg I’). We used Ξ2 -tests for categorical features and Kolmogorov-Smirnov tests for numerical ones, using a p-value cutoff of 0.05 for significance for both. Interval overlaps were calculated with bedtools (42).

### Comparison of exonized *Alu* elements between the GRCh38 and CHM13 genome assemblies

Annotations of genes and *Alu* repeat elements for the GRCh38 and the CHM13 v2.0 genome assemblies were obtained from the UCSC Genome Browser. *Alu* annotations of one genome were then projected onto the other using the ‘lift over’ tool, and ‘bedtools intersect’ was used to identify mappings to the local annotations. A match was determined if the features overlapped by >=25 bp.

### PCR Validation

#### Event selection

A total of nine gene-context (*i.e.*, intronic and in antisense to the gene) and one intergenic *Alu* sequences on human chromosome 12 predicted to be exonized by eXAlu were randomly selected from among those with tissue evidence from spliced alignments of GTEx RNA-seq data used for training and evaluation. Two examples did not render complete exon boundary predictions (inferred from spliced RNA-seq alignments) and hence were not pursued further, and initial primer design failed for two others. The primers are listed in **Supplemental Table S2**.

#### RT-PCR reaction

Commercially available total RNA isolated from Human Liver and Cerebellum was purchased (Ambion and Biochain) and converted into cDNA using the SuperScript II kit (Thermofisher) following the manufacturers’ recommended protocol. The resulting cDNA was amplified using the NEB Q5 High Fidelity Master Mix following the manufacturers recommended condition. Briefly an initial denaturation of 98C for 3 minutes was followed by 35 cycles of 98C for 10 second denaturation, 55-64C for 30 seconds for annealing followed by 72C for 30 seconds extension. A final extension at 72C for 2 minutes was performed following cycling.

### Detection and validation of candidate polymorphic *Alu* insertion-exonization events in the population

28,319 records of *Alu* insertion variants in the human genome previously reported in (27), predicted with the tool MELT (43) from whole genome sequencing data from the HapMap and 1000 Genomes Projects, were obtained from the International Genome Sample Resource portal (https://www.internationalgenome.org/). We generated the insertion and surrounding context sequences for 8,909 *Alu* retrotransposon insertions located in compatible GENCODE v.36 gene contexts (*i.e.*, intronic and in antisense to the gene) based on the VCF annotations, and ran our program eXAlu to predict instances of exonization. A mutated genome was generated, containing sequences for polymorphic *Alu* insertions predicted by eXAlu to be exonized, and RNA-seq reads were mapped using the spliced aligner STAR v.2.4.2a. Lastly, for each event we sought same-donor RNA-seq evidence using two approaches: *i)* read alignments were used to assemble transcripts using CLASS2 v.2.1.7, as before, and exons overlapping *Alu* insertions were selected for manual curation; and ii) spliced read alignments within the boundaries of the inserted *Alu* plus a 50 bp vicinity were similarly examined.

## ACKNOWLEDGMENTS

Program development and all calculations were performed on the ARCH Computing Center at the Johns Hopkins University and supported in part by instrumentation grant R01 GM129085-01SS. This work was supported in part by NIH award R01 GM129085 to L.F.. S.S. was supported in part by a grant from the Stanley Medical Foundation.

## CONFLICT OF INTEREST

The authors declare that they have no conflict of interest.

## AUTHORS’ CONTRIBUTIONS

L.F. and Z.H. conceived the project and computational methods and S.S. directed the validation experiments. Z.H. developed the model. Z.H., L.F., N.P. and G.P. tested the program and contributed to the computational analyses, and S.S. and O.C. performed the experiments. L.F., Z.H. and S.S. wrote the manuscript, and all co-authors read and agreed to the content.

## Notes

https://github.com/splicebox/eXAlu

### Competing Interest Statement

The authors have declared no competing interest.

