## Supplementary Tables and Figures for "Predicting *Alu* exonization in the human genome with a deep learning model"

SUPPLEMENTARY MATERIAL FOR THE ARTICLE:
"PREDICTING *ALU* EXONIZATION EVENTS IN THE HUMAN GENOME WITH DEEP LEARNING"
BY Z. HE, O. CHEN, N. PHILLIPS, G.I.M. PASQUESI, S. SABUNCIYAN AND L. FLOREA

**Table of contents**

**Supplementary Tables**

**Supplementary Table S1.** Distribution of data used for model training and testing.
**Supplementary Table S2.** List of PCR primers for experimental validation.

**Supplementary Figures**

**Supplementary Figure S1.** Selected predictions of *Alu* exonization events for validation with RT-PCR.
**Supplementary Figure S2.** Predicted exonization events at polymorphic *Alu* insertion sites and matching RNA-seq evidence from the same donor(s).

**Supplementary Table S1.** Distribution of data used for model training and testing. Raw RNA sequencing data was obtained from the GTEx repository and processed as described in the **Methods**.

| Tissue | #Samples | #*Alus* | #Unique *Alus* | #Exons | #Unique Exons |
|---|---|---|---|---|---|
| Ovary | 5 | 4952 | 2187 | 3703 | 1873 |
| Spinal_cord_cervical_c1 | 11 | 7140 | 2149 | 5269 | 1864 |
| Putamen_basal_ganglia | 11 | 7611 | 2419 | 5525 | 2081 |
| Breast | 12 | 10682 | 3366 | 7704 | 2866 |
| Substantia_nigra | 11 | 8039 | 2363 | 5891 | 2040 |
| Hypothalamus | 11 | 8487 | 2729 | 6258 | 2380 |
| Cerebellar_hemisphere | 11 | 12074 | 3359 | 9160 | 3087 |
| Lung | 4 | 3320 | 1871 | 2443 | 1526 |
| Hippocampus | 11 | 6950 | 2360 | 5093 | 2009 |
| Liver | 12 | 8595 | 2981 | 6120 | 2458 |
| Spleen | 5 | 3589 | 1782 | 2579 | 1422 |
| Colon | 13 | 10519 | 2936 | 7604 | 2531 |
| Frontal_cortex | 11 | 7468 | 2357 | 5525 | 2052 |
| Caudate_basal_ganglia | 11 | 8697 | 2681 | 6404 | 2345 |
| Adipose | 6 | 4501 | 2142 | 3283 | 1790 |
| Stomach | 6 | 1752 | 1129 | 1270 | 893 |
| Pancreas | 6 | 3691 | 1573 | 2596 | 1284 |
| Esophagus | 5 | 3271 | 1655 | 2394 | 1378 |
| Nucleus_accumbens_basal_ganglia | 11 | 9670 | 2925 | 7131 | 2582 |
| Cortex | 11 | 9113 | 2704 | 6770 | 2367 |
| Anterior_cingulate_cortex_BA24 | 11 | 6565 | 2230 | 4782 | 1884 |
| Small_intestine | 12 | 11541 | 3369 | 8342 | 2889 |
| Heart | 13 | 8228 | 2392 | 5892 | 2015 |
| Amygdala | 11 | 8083 | 2349 | 5923 | 2048 |
| Muscle | 12 | 8132 | 2409 | 5600 | 1974 |
| Cerebellum | 11 | 10432 | 3209 | 8011 | 2921 |
| Adrenal | 12 | 9041 | 2461 | 6541 | 2105 |
| Bladder | 10 | 9831 | 2951 | 7056 | 2530 |
| ALL | N/A | 211974 | 11930 | 154869 | 12206 |

**Supplementary Table S2.** List of PCR primers for experimental validation.

| Name | *Alu* exon coordinates | Junction | Primer sequence |
|---|---|---|---|
| chr12: 319_L56 | chr12:31,990,471-31,990,716 | L | AGGACACAGAAAGACAGCCA |
| chr12: 319_R206 | chr12:31,990,471-31,990,716 | R | TGGTTCACGCTACTCAGGAG |
| chr12: 319_L187 | chr12:31,990,471-31,990,716 | L | CTCCTGAGTAGCGTGAACCA |
| chr12: 319_R380 | chr12:31,990,471-31,990,716 | R | CAAATCCATCTGCACTGAAGC |
| chr12:505_L244 | chr12:50,605,842-50,605,964 | L | GCTCTCGGAGGATGGAGTTT |
| chr12:505_R441 | chr12:50,605,842-50,605,964 | R | TGCGGGCTGTAAGGAGATAG |
| chr12:505_L261 | chr12:50,605,842-50,605,964 | L | TTTCACTCTTGTTGCCCAGG |
| chr12:505_R439 | chr12:50,605,842-50,605,964 | R | CGGGCTGTAAGGAGATAGGAG |
| chr12:125_L37 | chr12:125,084,600-125,084,721 | L | GAGTGGTTCAAAGGCAGTCG |
| chr12:125_R225 | chr12:125,084,600-125,084,721 | R | GGGATACTGAAGTGGGAGGA |
| chr12:125_L151 | chr12:125,084,600-125,084,721 | L | AGTGCAGTGGTGTGATCTCA |
| chr12:125_R305 | chr12:125,084,600-125,084,721 | R | CAAAGCCACTTCTTGCCTCA |
| chr12:1215_R303 | chr12:121,554,120-121,554,230 | R | AGTGGGATCTGGCTCTGTTG |
| chr12:1217_L12 | chr12:121,782,883-121,782,947 | L | CGTTGTCGTAGCTGGTGGG |
| chr12:1217_L24 | chr12:121,782,883-121,782,947 | L | TGGTGGGATTCATGACGTCA |
| chr12:1217_R200 | chr12:121,782,883-121,782,947 | R | ACCCTGAACCTCTACGACAC |
| chr12:572_L133 | chr12:57,260,139-57,260,261 | L | CTGCATGACCACTACACCTG |
| chr12:572_R294 | chr12:57,260,139-57,260,261 | R | GGGCAGGGTCTCATTCTGT |

**Supplementary Figure S1.** Selected predictions of *Alu* exonization events for validation with RT-PCR. Examples at genes AC068987.2 and TTC41P were incomplete and were excluded from primer design and PCR validation.

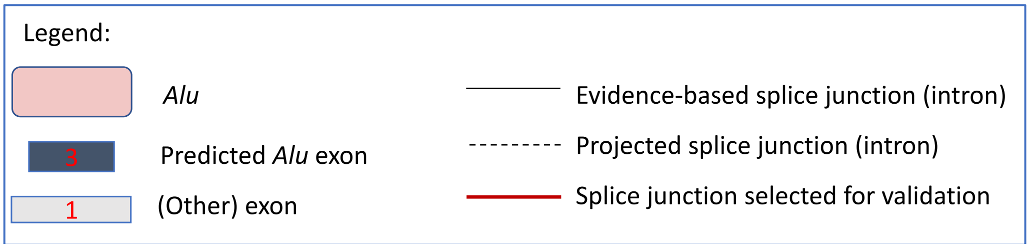


(1) #1: Alu: chr12:31990430-31990748 (-) *Alu*Sx ; RESF1 – Retroelement silencing factor 1 (+ strand, 5'->3')

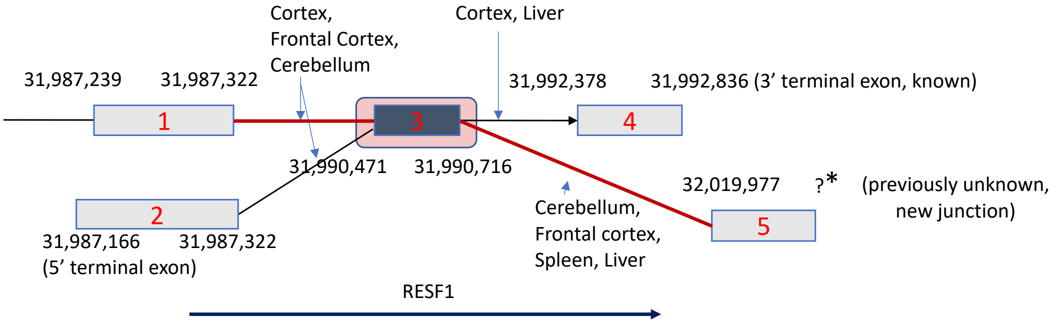


(2) #2: Alu: chr12:50,605,807-50,606,132 (-) *Alu*Sq ; DIP2B– Disco Interacting protein 2B Homolog (+ strand, 5'->3)

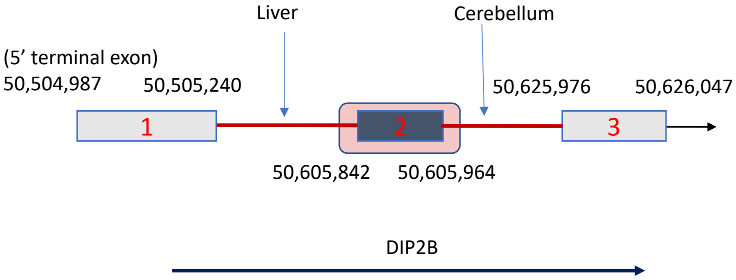


(3) #3: Alu: chr12:51,815,362-51,815,508 (-) *Alu*J ; AC068987.2 – (+ strand; 5'->3')

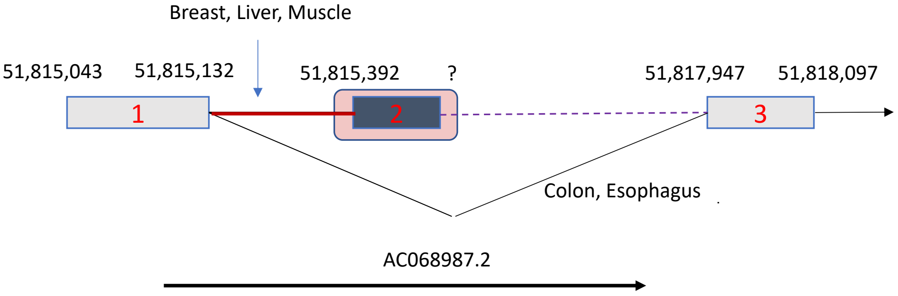

(4) #4: Alu: chr12:57,259,972-57,260,295 (+) *Alu*Jb ; R3HDM2- R3H Domain Containing 2 (- strand, 3’<-5’)

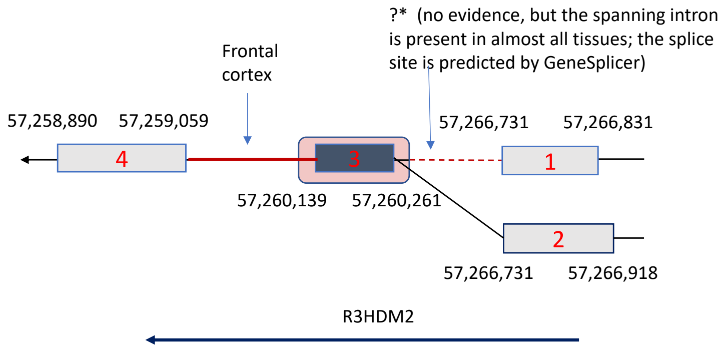


(5) #5: Alu: chr12:103,886,971-103,887,290 (+) *Alu*Sp ; TTC41P – Tetracopeptide Repeat Domain 41, Pseudogene

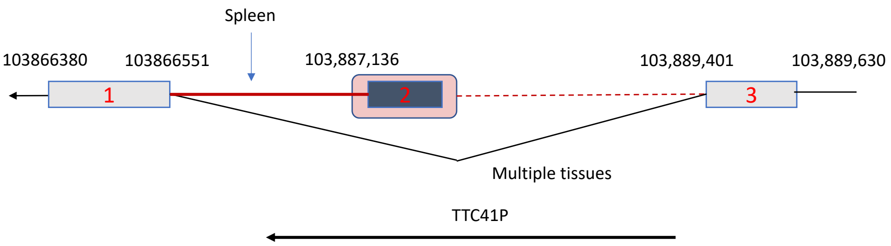


(6) #6: Alu: chr12:121,554,076-121,554,260 (+) *Alu*Jo ; KDM2B- Lysine Demethylase 2B (- strand, 3’<-5’)

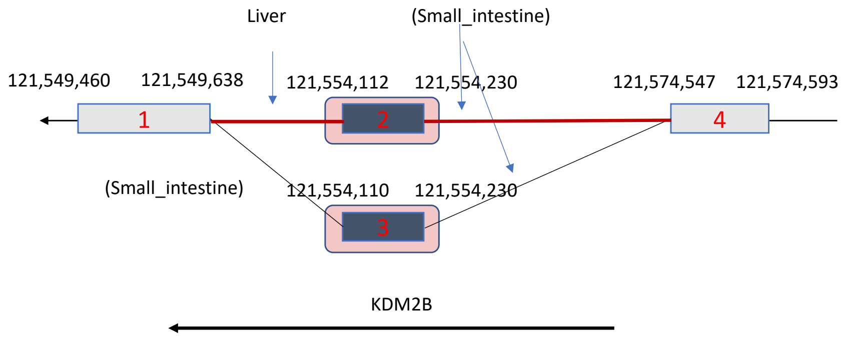


(7) #7: Alu: chr12:39,475,800-39,475,998 (-) *Alu*Jo ; Intergenic – no annotated gene/exon;
Alu exon boundaries based on GeneSplicer splice site predictions supported by evidence from flanking introns

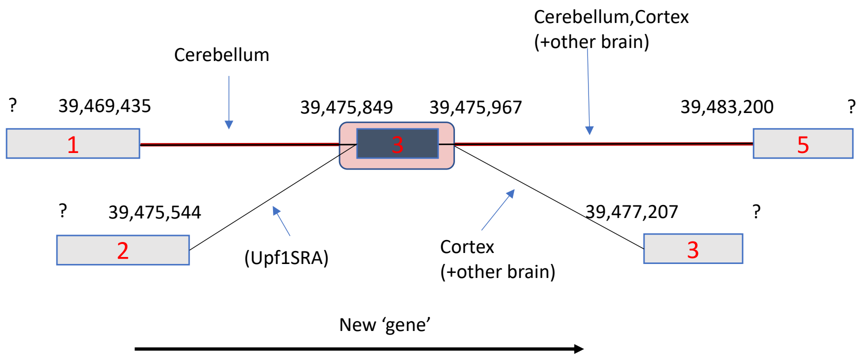

(8) #8: Alu: chr12:121,782,877-121,782,881 (+) *Alu*Jb ; RHOF- Ras Homolog Family Member F, Filopodia Assoc

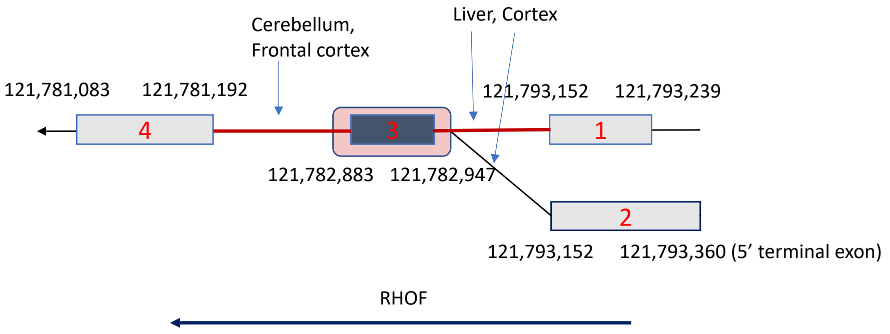


(9) #9: Alu: chr12:123,138,156-123,138,460 (+) *Alu*Jo ; PITPNM2 (- strand, 3'<-5')

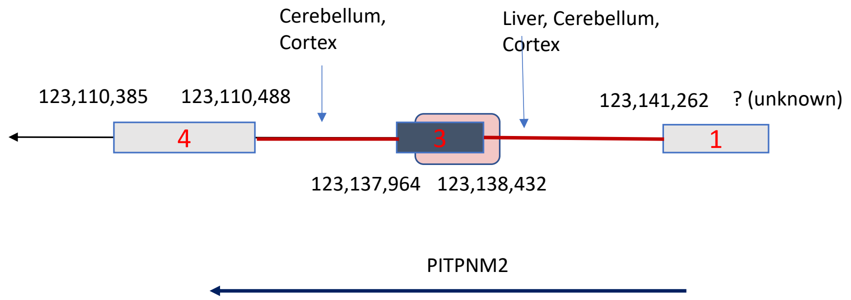


(10) #10: Alu: chr12:125,084,561-125,084,891 (-) *Alu*Jb ; AACS - Acetoacetyl-CoA Synthetase (+ strand, 5'->3')

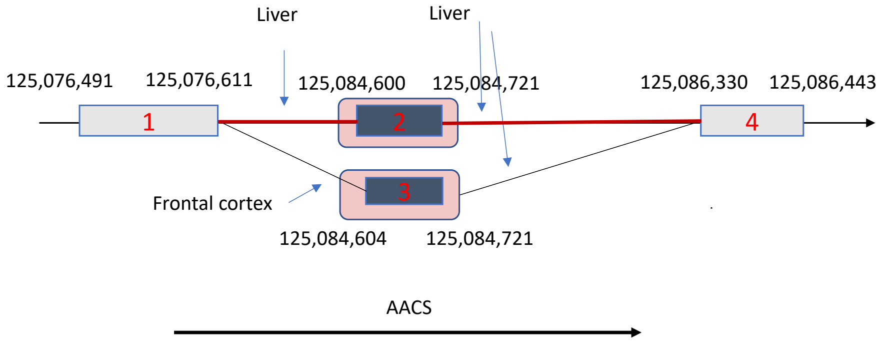

**Supplementary Figure S2.** Predicted exonization events at polymorphic *Alu* insertion sites and matching RNA-seq evidence from the same donor(s). (A, B) Potential polymorphic *Alu* exonization events at the EYA3 (A) and HMGCR (B) genes. In both cases, the *Alu* exon could potentially represent an alternative 5’ gene end; alternatively, evidence may be missing for the 5’ exon boundary. Numbers in parentheses represent the numbers of RNA-seq spliced read alignments supporting the splice junction. Coordinates are in the ‘mutated’ genome.

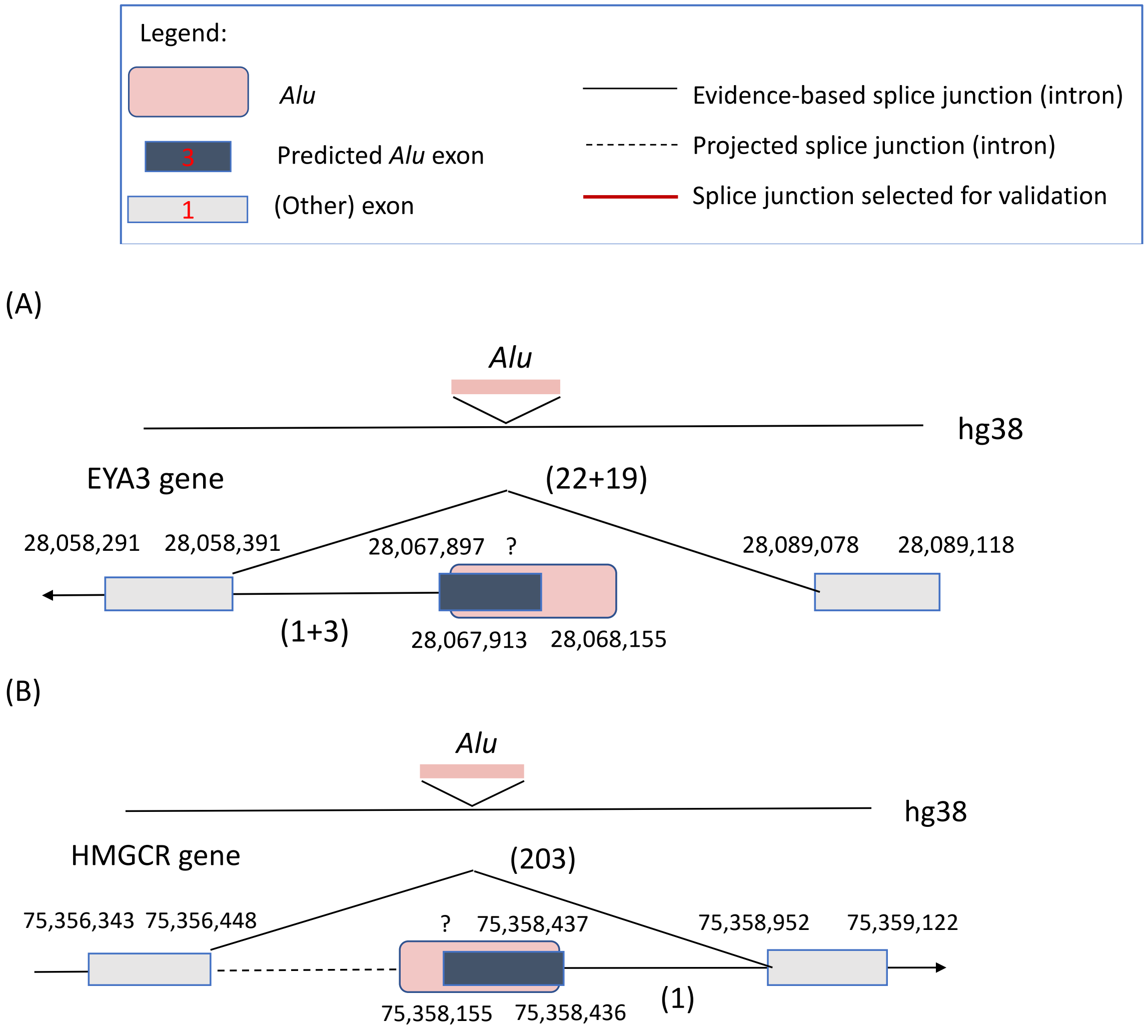
